# Reduced-observable dynamic mode decomposition for identifying theta and beta activity in working-memory EEG

**DOI:** 10.64898/2026.10.20.760227

**Authors:** David Chavez-Huerta, Mohammad Khosravi

## Abstract

The analysis of transient oscillatory activity in neural recordings remains challenging for linear methods, particularly when signals are nonlinear, noisy, and distributed across multiple channels. This paper introduces a Reduced-Observable Extended Dynamic Mode Decomposition (RO-eDMD) algorithm using a Random Fourier Features dictionary to characterize high-dimensional data in electroencephalography (EEG) recordings. The method extends the Koopman operator framework by incorporating rank reduction directly in the observable space and employing a randomized feature map that approximates the Gaussian kernel, enabling the capture of transient components without explicit dictionary optimization. To assess the method and demonstrate applicability, a logistic regression model is trained using a working memory paradigm as a test case, using dominant power values extracted via RO-eDMD in the theta band as a predictor and the type of event (encoding vs. fixation) as the dependent variable. We find correlations between oscillatory power and memory encoding events in the analyzed bands. Additionally, we find an increase in correlation associated with theta power and memory load, replicating similar results in the literature. The presented algorithm offers a computationally efficient, interpretable framework applied to a working memory paradigm dataset, but is relevant to the analysis of neural time series from diverse experimental contexts.

## 1 Introduction

Data-driven analysis of neural time series is fundamental to understanding brain function, yet the inherent nonlinearity, high dimensionality, and noisyness of electrophysiological recordings pose persistent analytical challenges. While linear methods such as principal component analysis, Fourier transforms, and autoregressive models have provided valuable insights, they often struggle to capture nonlinear dynamics and cross-channel interactions that are characteristic of cortical activity (Colbrook, 2023). This motivates the development of nonlinear, dictionary-based approaches that can reveal structure in complex neural recordings where linear methods reach their limits.

Among these approaches, Extended Dynamic Mode Decomposition (eDMD) (Williams et al., 2014) offers a promising framework. eDMD approximates the Koopman operator—a linear infinite-dimensional operator that governs the evolution of observables of a dynamical system—using a dictionary of nonlinear functions. By lifting the state space into a higher-dimensional observable space, eDMD can represent nonlinear dynamics in a linear form, enabling spectral analysis and reconstruction. However, several practical limitations hinder the application of standard eDMD to neural data: the computational cost of constructing and inverting the Gram matrix scales poorly with dictionary size; the choice of dictionary heavily influences performance yet remains largely heuristic; and reconstruction of transient signals, which are fundamental to neural activity, remains challenging.

In this paper, we introduce a Reduced-Observable Extended Dynamic Mode Decomposition (RO-eDMD) algorithm that addresses these limitations. This new method incorporates three key innovations. First, it achieves rank reduction directly in the observable space via singular value decomposition, avoiding explicit computation of the Gram matrix pseudoinverse and substantially reducing computational cost. Second, it employs a Random Fourier Features (RFF) dictionary (Rahimi and Recht, 2007, 2008) to approximate the Gaussian kernel, providing an efficient way to lift the state space. Third, the algorithm is shown to handle transient oscillatory signals (ubiquitous in neural recordings), which are notoriously difficult to reconstruct with Koopman-based methods.

We validate RO-eDMD on synthetic data with known transient oscillations and demonstrate its applicability to human electroencephalography (EEG) recordings from a working memory paradigm. This paradigm serves as an ideal test case as working memory is associated with well-characterized oscillatory activity in the theta and beta bands (Roux and Uhlhaas, 2014), providing a meaningful setting for evaluating whether the method can recover known signatures. While working memory provides the test case for this study, the method itself is general and applicable to a wide range of cognitive paradigms and neural recording modalities.

The use of working memory data also allows us to address frontal midline EEG activity during memory encoding and its correlation with memory load. Previous work (Onton et al., 2005) has identified frontal midline theta (fm*θ*) as a marker of cognitive effort, but the transient and trial-specific nature of this activity makes it difficult to characterize in detail with linear methods. The proposed RO-eDMD approach is specifically designed to capture such transient structure, enabling a more nuanced analysis of these phenomena.

The paper makes two primary contributions. First, we introduce an efficient and interpretable framework for modeling transient oscillatory activity in human EEG using RO-eDMD with Random Fourier Features: a methodological contribution that extends the Koopman operator literature by addressing practical implementation challenges. Second, we provide a test case of frontal midline dynamics associated with memory encoding, using logistic regression to link oscillatory power to event type and exploring the relationship between theta power and memory load.

The remainder of this paper is structured as follows: Section 2 introduces the notation; Section 3 presents preliminaries, including the working memory paradigm and an overview of DMD and eDMD; Section 4 describes the proposed RO-eDMD method in detail, including the Random Fourier Features dictionary and the algorithm structure in pseudocode; Section 5 presents results from synthetic validation and the working memory case study; Section 6 discusses our conclusions and future research directions.

## 2 Notation

Throughout this document, scalar quantities are written in lowercase characters, e.g., *n*; vectors are written in bold lowercase letters, e.g., ***x***; and matrices are written in bold, uppercase characters, e.g., **X**. Given a matrix **X***∈* ℝ^*I×J*^ , its column vectors are written as ***x***_*i*_ and its (*i, j*)-th scalar entry is denoted as *x*_*i*,*j*_. Operators and state spaces are written in uppercase calligraphic fonts, such as the Koopman operator K or the state space *ℳ*.

## 3 Preliminaries and related approaches

### 3.1 Paradigm Preliminaries

This article analyzes the data collected from a working memory paradigm with the analysis tools provided by RO-eDMD. Here, we briefly describe the paradigm for completeness. In (Onton et al., 2005), Onton investigates frontal midline activity during a visual task. The study focuses on the theta band, which has been previously linked to cognitive effort, heightened attention, and memory processes. The authors identify a set of EEG components that generate fmθ (frontal midline theta) activity. The study reveals that fmθ activity is not a single, uniform process but exhibits multiple spectral modes, including theta, low-beta (12–15 Hz), and delta (3 Hz) frequencies. The authors also analyze single-trial variability, demonstrating that the observed mean spectral changes are driven by a small proportion of trials with particularly strong theta activity. The findings suggest that fmθ dynamics may reflect dynamic adjustments in the medial frontal cortex to trial-specific behavioral contexts and task demands. The experimental paradigm used in the study was a modified version of the Sternberg task. Participants were presented with a series of eight consonant letters, some of which were to be memorized and others to be ignored. The letters were presented sequentially with a fixed stimulus onset asynchrony (SOA) of 1.4 seconds, and their duration was either 1.2 seconds (long condition) or 0.15 seconds (short condition). After the letter series, there was a 2–4 second maintenance period during which participants rehearsed the memorized letters. A probe letter then appeared, prompting participants to indicate whether it was part of the memorized set. The task was designed to manipulate memory load (3, 5, or 7 letters to be memorized) and to examine how EEG activity varied with increasing cognitive demand.

### 3.2 Dynamic mode decomposition (DMD) and extended dynamic mode decomposition (eDMD)

Dynamic mode decomposition (DMD) is a data-driven method for analyzing high-dimensional dynamics by decomposing them into *spatial modes* and corresponding temporal frequencies (Brunton et al., 2014). In the context of EEG analysis, DMD aims to identify recurring spatiotemporal patterns: oscillatory activity across electrodes that evolve in time with a specific frequency. Consider a neural activity recording (discrete data) consisting of *n* channels (dimensions) and *m* snapshots (data points), sampled at a constant interval Δ*t*. For each *k ∈* {1, … , *m*} , snapshots from the time instant *k*Δ*t* are arranged in vector columns ***x***_***k***_ *∈* ℝ^*n*^ of the data matrices **X, Y** *∈* ℝ^*n×*(*m−*1)^

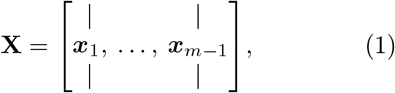

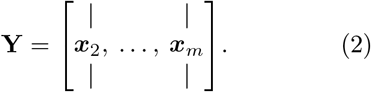

For our purposes, **Y** is a version of **X** *advanced one step in time*. DMD generates a matrix **A** that approximates how one brain activity state evolves into the next one, i.e., **AX** ≈ **Y**, assuming that the next state can be approximated from the current state using a linear transformation. The algorithm depends on the singular value decomposition (SVD) of **X**, given as

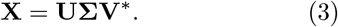

where **U** and **V** are unitary matrices, and **Σ** is a rectangular diagonal matrix with non-negative real entries. Defining matrix **A** := **U**^*\**^**YVΣ**^*−*1^ and calculating its eigendecomposition **AW** = **WΔ**, the original data **X** can be reconstructed by

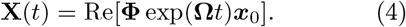

The matrix of eigenmodes **Φ** is defined as **Φ** ≜ **X**^*′*^**VΣ**^*−*1^**W** and continuous time eigenvalues are given by **Ω** = log(**Λ**)*/*Δ*t*.

The eigenvalues obtained from this decomposition encode essential dynamical information: their magnitudes indicate the level of stability (i.e., whether oscillations grow, decay, or persist), while their imaginary parts determine oscillation frequencies. The modes **Φ** represent spatial activation patterns across electrodes, revealing which brain regions participate in each oscillatory component. Together, the eigenvalues and modes provide a compact description of the spatiotemporal structure of the neural data.

Even in noiseless datasets, the DMD algorithm struggles to reconstruct transient signals and traveling waves (Brunton et al., 2014). This limitation appears because standard DMD assumes a single approximately linear evolution and relatively stable oscillatory behavior throughout the entire recording. However, neural signals often contain sudden changes, nonlinear effects, and nonstationary dynamics; such features violate that assumption. A possible solution is to apply the algorithm to small, overlapping data intervals known as mrDMD (Kutz et al., 2015). Besides a significant increase in computational cost, this partly defeats the purpose of generating a dynamical representation, as every analyzed window contains approximately linear signals with little relation to the whole data.

Extended dynamic mode decomposition (eDMD) is an extension of DMD (Williams et al., 2014) that addresses some of these limitations. Rather than analyzing the raw state directly, eDMD transforms the data into a feature space using a dictionary of observables—a set of nonlinear functions of the state—to produce more accurate approximations of the underlying dynamics. This lifting to a higher-dimensional space allows the method to better capture nonlinear dynamics that would be invisible to standard DMD.

Assume that the system evolution is defined by a corresponding discrete-time dynamical system (ℳ , *k, ℱ*), with ℳ *⊆* ℝ^*n*^ the state space, *k ∈ Z* the discrete time, and ℱ : ℳ → ℳa forward operator defined by its action on the elements of the state space, namely **x**_*k*+1_ = ℱ (**x**_*k*_). Given functions (observables) of the state space *ψ* :ℳ → ℂ, the Koopman operator has the defining property

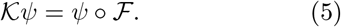

In other words, the Koopman operator *K shifts by one step in time* both functions of individual states ***x***_*k*_ and functions of the entire data set **X**. More precisely, for any *k ∈* ℤ, we have

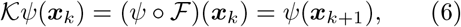

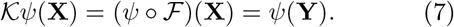

The approximation to the Koopman operator is given as K = **G**^*†*^**A**, where

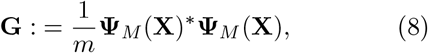

is the Gram matrix and

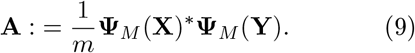

With **G**^*†*^ the conjugate transpose of **G**.This formulation has a key advantage: while standard DMD can only lift dynamics into a space of linear functions, eDMD can choose nonlinear observables to identify dynamical evolution in the observable space (Williams et al., 2014). By choosing an appropriate dictionary—a topic we address in Section 4.2—the method can capture complex neural phenomena such as transient oscillatory bursts.

In order to reconstruct states in ℳ, eDMD requires tuples of eigenvalues, eigenfunctions, and modes (*λ*_*i*_, *Φ*_*i*_, ***ν***_*i*_) associated with this finite approximation to *K*. The starting point of this algorithm is the work presented in (Williams et al., 2014). We propose a Reduced-Observable version that is

1. Focused on efficient computation of the Koopman operator, taking advantage of the SVD rank-reduction directly in the observable space, so that **G**^*†*^ is not explicitly calculated and inversion operations are scarce and performed on reduced matrices.
2. Comprehensive. Explicitly mentioning details of the algorithm that are often omitted in eDMD descriptions (such as Hankel-stacking, rank reduction, eigenvector projection, and dictionary construction) that are unavoidable for implementation.

This allows for a straightforward implementation in the context of neural time series. The reconstruction of **X** relies on the iterative application of *K*, so instead of generating ***x***_*i*+1_ from ***x***_*i*_; the complete matrix **X** is generated from the initial conditions vector ***x***_0_.

## 4 Methods

### 4.1 Proposed algorithm: RO-eDMD

In practice, the data matrices **X** and **Y** are structured as Hankel matrices before applying eDMD, where copies of the data are stacked vertically. Given *h ∈*ℕ, the Hankel data matrix **X** is defined as

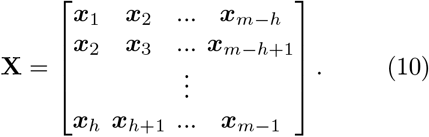

For simplicity. no additional notation is introduced to denote the stacked matrices **X** and **Y**. Both now have *hn* rows and *m*− *h* columns, but we choose to keep the variables *n* and *m* to denote their dimensions, as it makes the notation cleaner. Stacking parameter *h* must be chosen at the very least as *h* ≥ 2, as this warrants linear consistency in *X* and allows DMD and eDMD to encode standing waves (Tu et al., 2013). Literature recommends (Brunton et al., 2014) choosing the smallest integer *h* such that *hn >* 2*m*, where *n* is the number of channels and *m* is the number of snapshots in the data matrix.

In the coming sections, we implement the analysis with *h* = 5 to comfortably fulfill the previous guideline. Note that the effective number of channels is given by *d* · *m* and that the choice of parameter *h* is a trade-off between capturing temporal dynamics and maintaining computational efficiency. Next, we enumerate the steps of the algorithm.

1. Define a dictionary of *d* complex-valued observables D = {*ψ*_1_, … *ψ*_*d*_} and a vector-valued function ***ψ*** as (11). Subsequently, an associated matrix-valued function **Ψ**_*M*_ : ℝ^*n×m*^ → ℂ^*m×d*^ as (12)

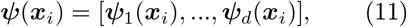

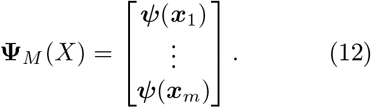 For **x**_1_, … , **x**_*m*_*∈*ℝ^*n*^, the appropriate choice of observables for the dictionary is, in general, an open problem. Our choice is discussed in sub-section 4.2. Hermite polynomials, radial basis functions, and discontinuous spectral elements are commonly used as dictionaries. Ideally, the properties of the dictionary set should reflect the dynamics of the system.
2. In this methodology, the SVD is computed and only the dominant features are retained, enabling more efficient computation: Compute the singular value decomposition (SVD) of the observables matrix 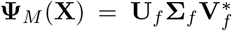.The Eckart-Young-Mirsky theorem warrants that the best approximation of **Ψ**_*M*_ (**X**) with maximum rank *r* is

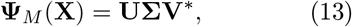

where matrices **U, Σ** and **V** are built with the first *r* columns ^1^ of **U**_*f*_ , **Σ**_*f*_ and **V**_*f*_ , respectively.
3. A finite approximation to the Koopman operator 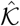 is to be calculated using the observable function matrix **Ψ**_*M*_ (**Y**). Let **G** and **A** be respectively given in terms of the SVD decomposition of **X**:

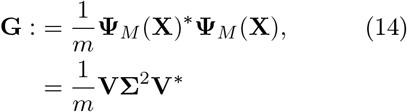

and

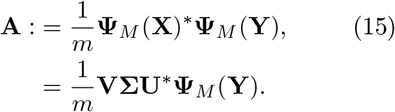 Also, define the Moore-Penrose pseudoinverse of **G** as **G**^*†*^ = **VΣ**^*−*2^**V**^*\**^. Then, the finite approximation of the Koopman operator K = **G**^*†*^**A** can be obtained as

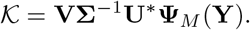 Note that there is no need to actually calculate **G**^*†*^. Through a similarity transformation 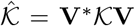, one can make the dimensions of the Koopman operator coincide with the rank reduction carried from the decomposition of **Ψ**_**M**_(**X**). Calculate the rank-reduced Koopman operator approximation as

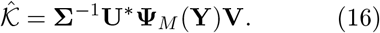 The implementation of SVD on the observable function matrix allows for a more efficient algorithm by avoiding the direct calculation of **G**^*†*^, through a similar way to the kernel approach in (Williams et al., 2014).
4. Compute the eigendecomposition of 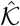 for both right and left eigenvectors, respectively

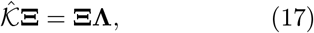

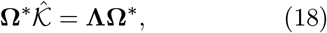

where **Ξ** is the matrix of right eigenvectors ***ξ***_*i*_ for *i* = 1, … , *r*, **Λ** is a diagonal matrix containing the *r* eigenvalues *λ*_*i*_, and **Ω** is the matrix of left eigenvectors ***ω***_*i*_, for *i* = 1, … , *r*. We scale **Ω** such that we have **diag**(**ΩΞ**) = *I*.
5. For a given ***x***_*i*_ in the state space ℳ , we define ^2^ the eigenfunctions of 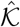 as *Φ*_*k*_ = ***ψ***(***x***_***i***_)**V***ξ*_*k*_. A diagonal matrix of eigenfunctions **Φ** is defined as

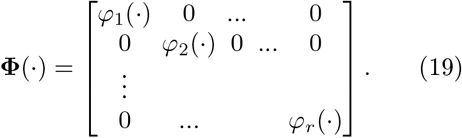
6. Define **Λ**_*M*_ as a matrix of eigenvalues of 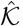, incorporating the powers of *λ*_*i*_ needed to generate *m* time steps in the algorithm implementation, i.e.

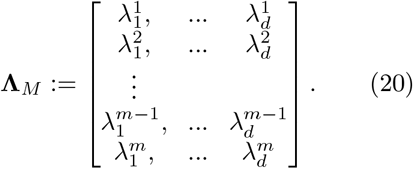
7. A weight matrix **B** needs to be built, such that a *full state observable* function *g*(**X**) = **X** can be defined as

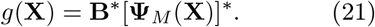 Using the already available SVD decompostion, calcualte **B** as

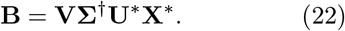
8. Compute the eDMD modes 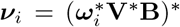. Subsequently, define matrix **V** as

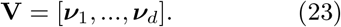
9. The algorithm operates in an observable (feature) space, but the final goal is to reconstruct the original data **X**. For the initial conditions vector ***x***_1_, the discrete-time dynamics reconstruction in matrix form **X**_r_ is generated by a forward operator *ℱ*_*M*_ defined as

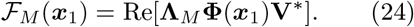 As a consequence of the Hankel stacking, *ℱ* _*M*_ generates *h* copies of the reconstruction

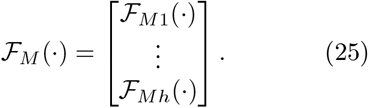

Since the Hankel matrix contains *h* shifted copies of the same time series, the reconstruction yields *h* versions of the dynamics, one for each shift. These copies are redundant; we retain only the first block ℱ_*M*1_(·), to avoid duplicating information and to ensure that the reconstructed matrix **X**_*r*_ has the same dimensions as the original data **X**:

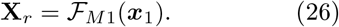

By means of the algorithm, **X**_r_ will approximate **Y**.

The proposed modifications to the standard eDMD algorithm are not merely algebraic; they significantly reduce computational cost and improve numerical stability in high-dimensional observable spaces. The accuracy of the reconstruction crucially depends on the elements of the dictionary *D* and its potential to reflect the dynamics. This is particularly important when reconstructing nonlinear phenomena. The next section describes the dictionary chosen in this paper.

### 4.2 Random Fourier Features Dictionary

RFFs are kernel approximations introduced in (Rahimi and Recht, 2007, 2008). Consider a positive definite shift-invariant kernel function *k*(·, ·) : ℝ^*n*^ × ℝ^*n*^ → ℝ with

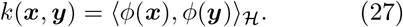

where *ϕ* : ℝ^*n*^ → *ℋ* is a suitably chosen *feature map*, lifting state vectors to Hilbert space *ℋ* equipped with inner product *⟨*·,·*⟩* _*ℋ*_. Instead of evaluating *k*(***x, y***) for all data points (***x, y***), RFF approximates the kernel as

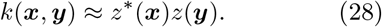

where *z* : ℝ^*n*^ → ℝ^*D*^ is a randomized feature map. An option for the RFFs mapping *z* is

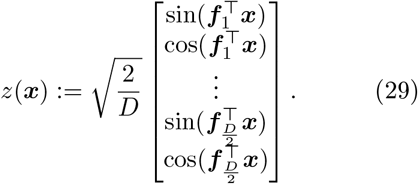

where **f**_*i*_ *∈* ℝ^*n*^ are sampled from a Gaussian distribution 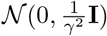, with **I** the identity matrix of appropriate dimensions. A more common *z* mapping disregards the sine terms and adopts a random frequency shift added to the cosine term. In (Sutherland and Schneider, 2015), it is shown that the sine-cosine version in (29) reduces several measures of error and thus, it is employed in the approach proposed here.

The choice of the Gaussian distribution for sampling the frequency vectors ***f***_*i*_ is motivated by Bochner’s theorem, which states that any shift-invariant kernel can be represented as the Fourier transform of a probability measure. For the Gaussian kernel, defined as:

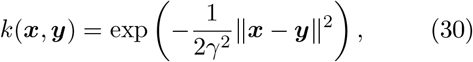

also known in the literature as the radial basis function (RBF) kernel, the inverse Fourier transform yields the sampling distribution:

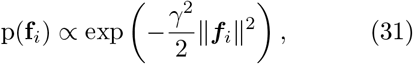

which corresponds precisely to 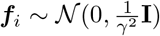. The parameter *γ* is a kernel bandwidth parameter that influences the spread of the frequency distribution and the smoothness of the resulting kernel approximation:

- When *γ* is large, frequencies are sampled with smaller variance, yielding less high-frequency components.
- When *γ* is small, frequencies are sampled with greater variance, yielding more high-frequency components.

For the analysis in this manuscript, we set *γ* = 200 as a default value, which works well for normalized input data. This choice was made for simplicity and to demonstrate the method’s general applicability, though we note that several optimization strategies can be deployed to systematically select *γ*.

Sampling from the Gaussian distribution concentrates samples near the frequencies that matter most for the signals of interest. In contrast, sampling frequencies from a uniform distribution would inevitably allocate samples to irrelevant frequencies (Rahimi and Recht, 2007), requiring substantially more samples to achieve the same approximation quality, since it fails to exploit the spectral properties of the system dynamics. This is particularly relevant to our application, since EEG signals of interest are typically contained in specific frequency bands, making Gaussian sampling a natural and efficient choice.

In our algorithm, we use the mapping in (29) to generate a matrix of observables **Ψ**_*M*_ (*X*). More precisely, a set of 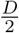 vectors ***f***_*i*_ are stacked in the matrix **F** *∈* ℝ^*D×n*^ as

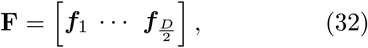

and then, the matrix of observables **Ψ**_*M*_ (*X*) ℝ^*m×D*^ is constructed as

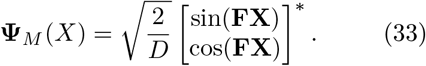

### 4.3 Frequency decomposition and power

Beyond the reconstruction of **X** from the vector of initial conditions ***x***_1_, eDMD facilitates the analysis of the system on the tuple (*λ*_*i*_, *Φ*_*i*_, ***ν***_*i*_). In particular, the frequencies associated with the Koopman modes can be calculated as:

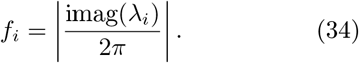

The power associated with them is given by the eigenfunctions *Φ*_*i*_:

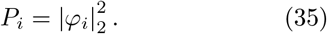

Once repeated and negligible values are omitted, the relation of power *P*_*i*_ and frequency *f*_*i*_ serves as a direct view of the dataset frequency content, akin to FFT-based techniques.

### 4.4 Algorithm Structure

For clarity, the key steps of the RO-eDMD algorithm with RFFs are summarized below in pseudocode form. MATLAB code is available in a GitHub repository (Chavez-Huerta, 2026).

#### Algorithm 1 RO-eDMD

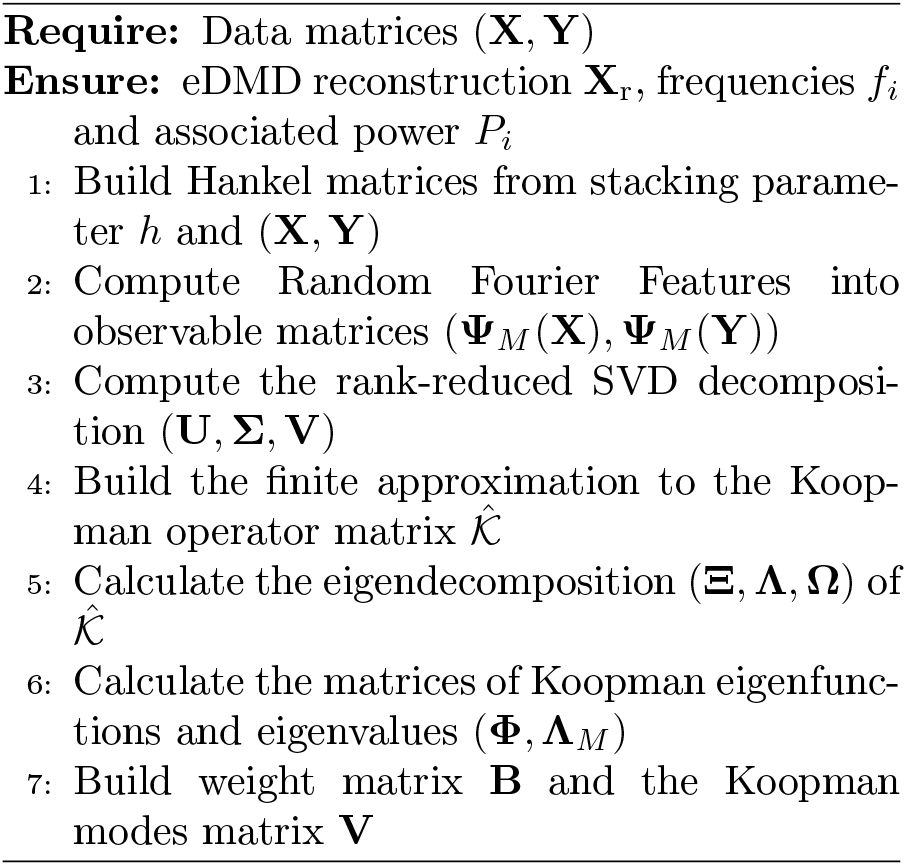

### 4.5 Synthetic dataset and EEG data

A synthetic dataset is constructed and tested in MATLAB R2024b to demonstrate the algorithm and validate its ability to recover transient oscillatory signals embedded in noise. The dataset consists of a 40-channel time series with 2000 snapshots and contains transient oscillations at 68, 21, 35, and 9 Hz. The noise signal is additive zero-mean white Gaussian noise (AWGN) generated with MATLAB’s randn function. Different levels were obtained by scaling the generated noise with a constant coefficient, resulting in datasets with measured signal to noise ratios (SNRs) of approximately 41 dB, 30.02 dB and 6.79 dB. The synthetic datasets are shown in Fig. 1. All figures in the manuscript were generated using MATLAB.

**Fig. 1.**
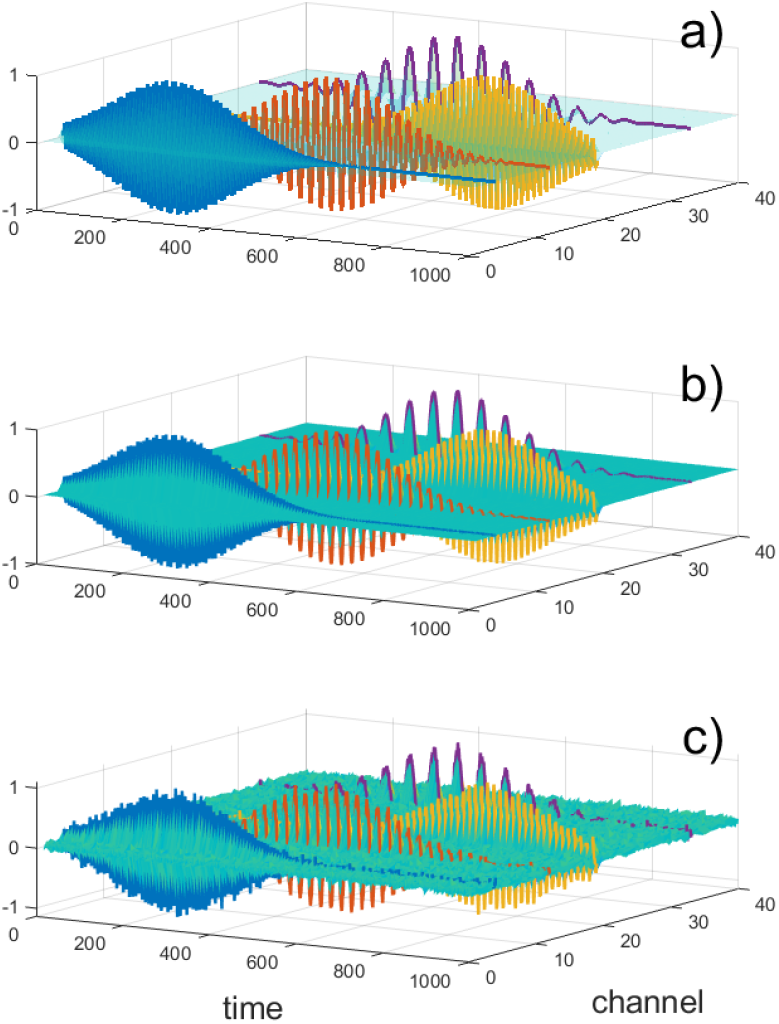
Synthetic dataset. Transient sinusoids in a 40-channel array. In a) a nearly noise-free (41 dB SNR) dataset, in b) 30.02 dB and c) 6.79 dB SNRs.

Additionally, the EEG data generated in the working memory paradigm described in Section 3.1 is analyzed to validate the algorithm. The data was obtained from the OpenNeuro database (Onton et al., 2022). The data accession number is ds004117. From the 71 original channels, a subset is selected consistent with the anatomical correlations of scalp EEG described in (Koessler et al., 2009). The resulting frontal midline cluster includes channels Fz, FCz, Cz, CPz, AFz, F1, and F2, which are investigated for memory-related activity.

Data from all 23 participants are processed. For each subject, epochs corresponding to encoding events are extracted and concatenated. We consider encoding events as EEG data from the frontal midline cluster when a character is presented as the task role is *to remember*. In particular, we only process events from trials where at least five letters are presented with this role. Equivalent-length fixation periods, during which no letters are displayed, serve as control events. For subjects 14 to 24, the data are downsampled to 250 Hz to ensure consistent sampling rates across all subjects before applying the algorithm.

A band-pass filter of 0.01 to 100 Hz has already been applied to the dataset and no further pre-processing is carried out. Once preprocessed, every valid encoding event from every participant is subjected independently to the algorithm. While our preprocessing approach differs from the original study by Onton et al. (2005), who employed ICA-based component decomposition and artifact rejection, our simpler pipeline was deliberately chosen for three reasons. First, in Delorme (2023) it is demonstrated that additional preprocessing steps such as re-referencing, baseline removal, and ICA-based artifact correction do not necessarily improve statistical sensitivity in EEG analyses. Second, our RO-eDMD algorithm operates directly on channel data and already incorporates a nonlinear dictionary that can capture structure without requiring ICA-based source separation. Third, the RFF dictionary’s ability to approximate kernel methods provides a degree of robustness to noise and artifacts that parallels the benefits of ICA decomposition. These differences in preprocessing affect direct comparability of results; however, our primary interest is in demonstrating the utility of RO-eDMD as an analytical tool rather than precisely replicating the original study’s findings.

### 4.6 Validation Strategy

To evaluate the proposed RO-eDMD algorithm, we employ a two-pronged validation approach that assesses both reconstruction fidelity and spectral recovery accuracy in an artificial dataset. The use of an artificial dataset makes the validation straightforwards, as the ground truth is available.

First, we quantify reconstruction accuracy using the Normalized Root Mean Square Error (NRMSE) between the original data **X** and the reconstructed data **X**_r_:

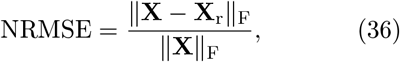

where ∥ · ∥_F_ denotes the Frobenius norm. This metric provides a global measure of how well the algorithm reproduces the original time series. To test the algorithm’s robustness, we systematically corrupt the data with additive white Gaussian noise at varying Signal-to-Noise Ratio (SNR) levels. This allows us to characterize the algorithm’s performance degradation under realistic recording conditions.

Second, we assess spectral recovery accuracy by analyzing the dominant frequencies extracted from the eigenvalues *λ*_*i*_ of the Koopman operator, as the eigenvalues encode the oscillatory frequencies of the identified modes. By comparing the recovered dominant frequencies against the true frequencies used to generate the synthetic data, we can determine the algorithm’s ability to correctly identify the underlying dynamics.

To evaluate the RO-eDMD algorithm against established Koopman-based approaches, we compared its performance against a standard DMD algorithm and an eDMD algorithm over the same synthetic dataset. All three methods were applied to the 40-channel time series across three noise levels. For eDMD, we employ a second order polynomial dictionary and the Hankel stacking parameter h = 5. The small polynomial order of the dictionary allows compute times of around 60 seconds.

## 5 Results

### 5.1 Synthetic dataset implementation

Given the initial conditions provided by the first data snapshot, RO-eDMD reconstructions are plotted in Fig. 2. Transient signals are inherently challenging to reconstruct using Koopman-based approaches because the Koopman operator assumes linear evolution in the observable space, as is the case for signals with persistent oscillatory dynamics. We deliberately choose to present these challenging transient signals because they are fundamental to neural activity and represent a realistic test case for the proposed method. For completeness, we note that RO-eDMD reconstructs non-transient oscillatory signals (with constant, growing, and decaying amplitudes) with excellent accuracy; however, we do not focus on these results since this task is already established in the literature for standard eDMD, and our present work focuses on the transient case.

**Fig. 2.**
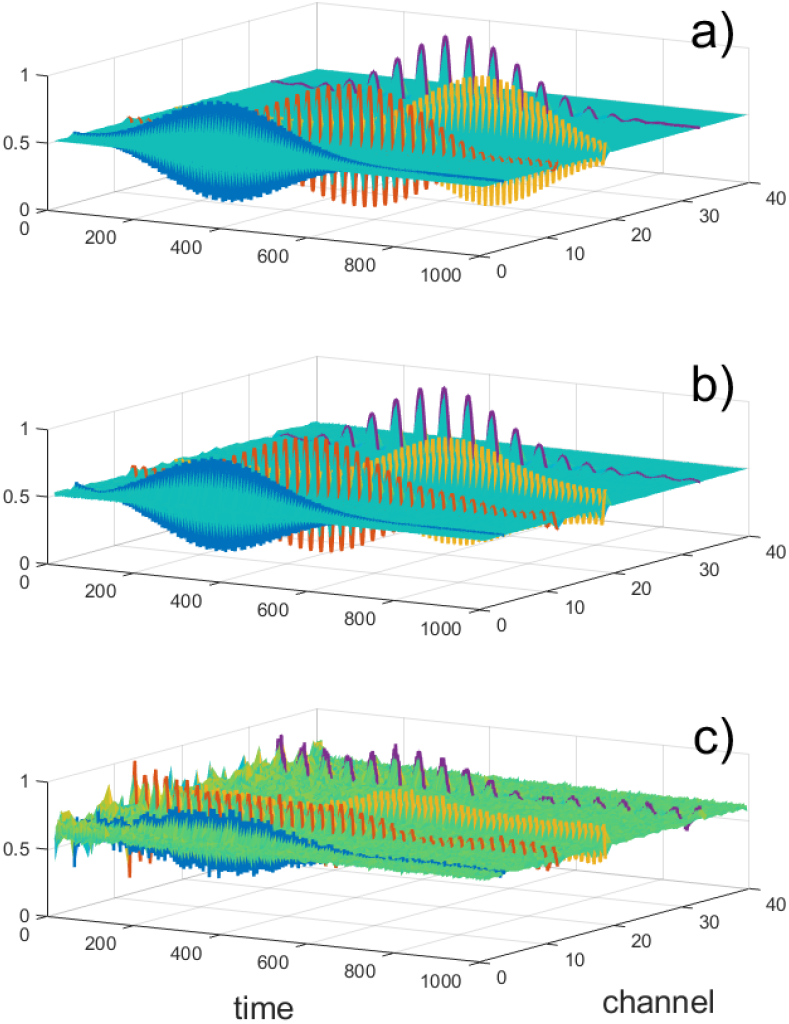
RO-eDMD reconstruction for SNRs: 41 dB, 30.02 dB and 6.79 dB, respectively

If the SNR is significantly reduced, the characteristics of the signal become harder for the algorithm to recover. In particular, transient signals can be shifted, deformed, or suppressed as noise increases. Nevertheless, frequency contents are still retrievable for many noise scenarios.

We calculate the NRMSE for the three noise levels used before and plot them in Fig. 3, it is higher in the most noisy dataset, yet is very similar in the 41 dB and 30.02 dB cases.

**Fig. 3.**
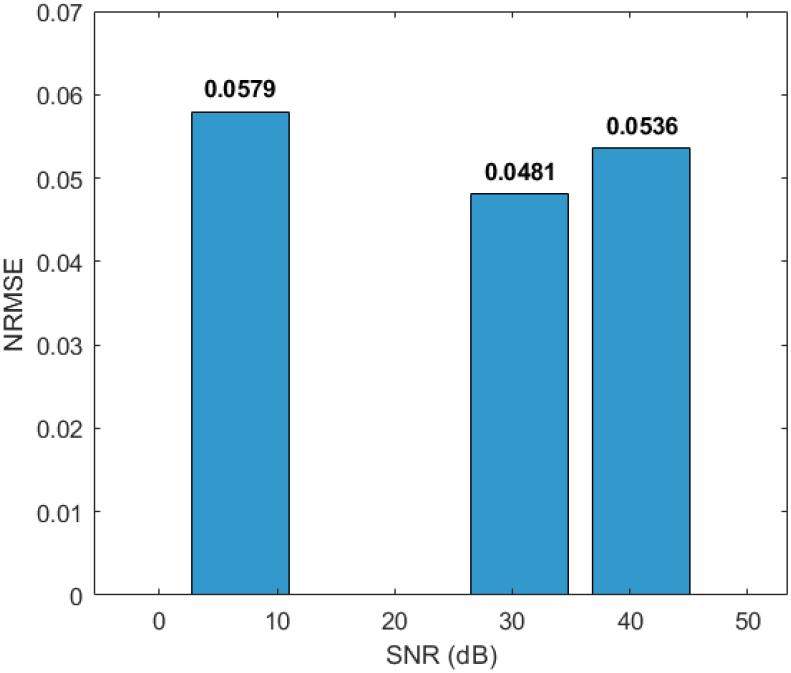
NRMSE values per SNRs. Model accuracy is reduced at the lowest tested SNR. At 41 dB and 30.02 dB, accuracy is similar.

A power to frequency plot is presented in Fig. 4, calculated as in Eqs. (34) and (35). The frequency of the two main signals in the synthetic dataset are registered as dominant peaks in the vicinity of 68, 21, 35 and 9 Hz.

**Fig. 4.**
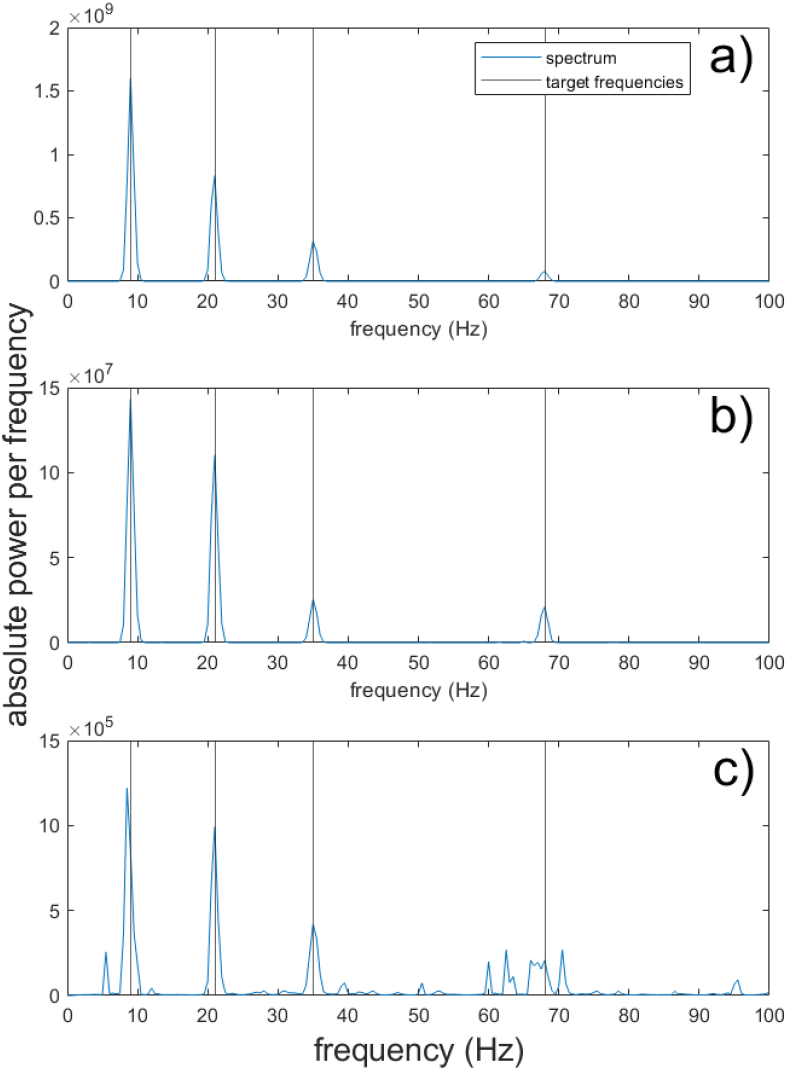
Power *P*_*i*_ values plotted against their corresponding frequencies *f*_*i*_ *∈* [0.1, 100] Hz. Frequencies in the original synthetic dataset are marked as black vertical lines. SNRs of 41 dB, 30.02 dB and 6.79 dB, plotted in a), b) and c), respectively.

As power *P*_*i*_ is a function of the right eigenvectors ***ξ***_*k*_, only their relative magnitude is meaningful and they can be normalized. Koopman-based methods can be sensitive to parametrization in unexpected ways. Currently, there is no general consensus on several aspects of dictionary design, rank truncation and parameter optimization (Colbrook, 2023) and some trial and error can be necessary to set up the algorithm.

In Fig. 5, we show a comparison of standard DMD, an eDMD algorithm, and RO-eDMD over the same synthetic dataset at the three noise levels. DMD reproduces oscillations but is not able to capture it’s transient nature or accurately identify it’s frequencies. eDMD over a polynomial dictionary of second order is computationally expensive (see Table 1), struggles to capture transient behavior, and correctly identifies some frequen-cies. RO-eDMD shows better performance both at transient reconstruction and frequency identification. All reconstructions are phase shifted to account for Hankel-stacking before calculating NRMSE. Frequency, phase and transient inaccuracy result in higher NRMSE values for both DMD and eDMD decompositions. Average compute times are recorded in Table 1, as a measure of performance in relation to user experience.

**Table 1.**
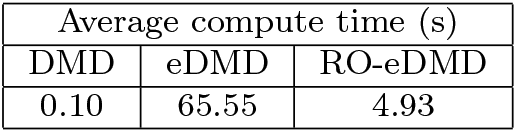
Average compute times on the artificial dataset.

| Average compute time (s) |  |  |
| --- | --- | --- |
| DMD | eDMD | RO-eDMD |
| 0.10 | 65.55 | 4.93 |

**Fig. 5.**
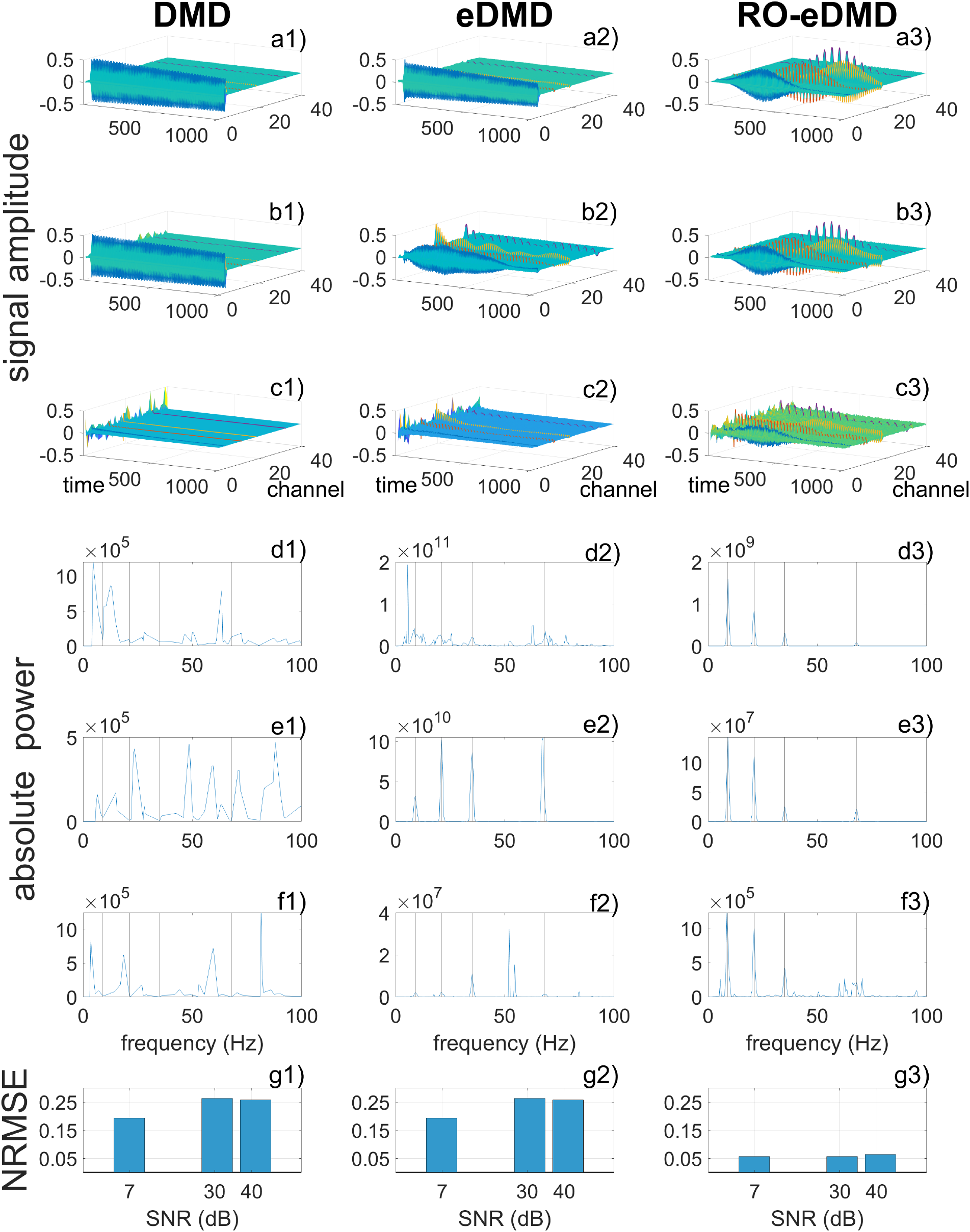
Performance comparison of Koopman-based methods on over the artificial dataset. The figure evaluates three algorithms: standard DMD (left column), eDMD over a polynomial dictionary (center column), and RO-eDMD over a RFF dictionary (right column) across decreasing SNRs. (a-c) Surface plots of the reconstructed data. From top to bottom, SNRs are 41 dB, 30.02 dB and 6.79 dB. (d-f) Corresponding power spectra obtained from each algorithm’s eigenvalue decomposition. Vertical dashed lines indicate the four true transient frequencies (9, 21, 35, and 68 Hz). From top to bottom, SNRs are 41 dB, 30.02 dB and 6.79 dB. (g) Bar plots comparing NRMSE of the full reconstructed signal against the original signal for each algorithm.

### 5.2 Implementation on a working memory paradigm

Fig. 6 presents the reconstruction of a letter-encoding event in the electrodes of the frontal midline cluster during 350 snapshots (encompassing 1.4 s). The reconstruction displays a burst of activity in the channels rendered at the front of the 3D plot, corresponding to Fz, FCz, and Cz. Additionally, notice a small collection of impulse-like transients in the background channel (corresponding to F2) reconstructed with a noticeable shift towards the beginning of the dataset.

**Fig. 6.**
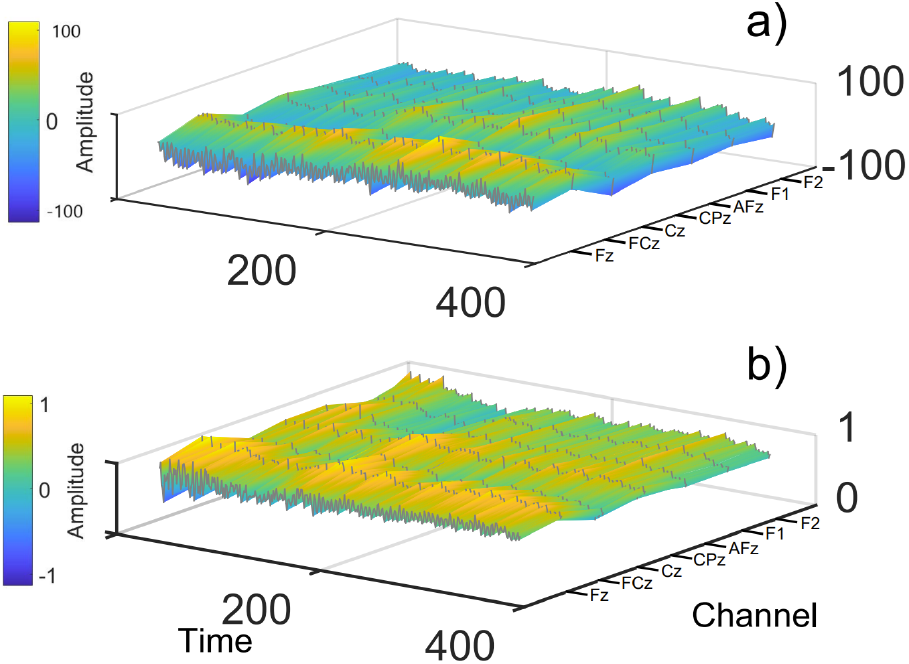
a) Activity in the frontal midline cluster during a particular encoding event. b), reconstructed and normalized data produced by the RO-eDMD algorithm. Time is shown on the x-axis, channel index on the y-axis, and signal amplitude is displayed both on the z-axis and the plot coloring, as indicated in the color bar. Note how transient characteristics of the signal are shifted in time, specially in the Fz and FCz channels.

Eigenvalues *λ*_*i*_ are used to derive the oscillatory frequencies associated with each mode, as defined in (34). The frequencies are grouped into the electrophysiological bands: *delta* (0.5–4 Hz), *theta* (4–8 Hz), *alpha* (8–12 Hz), *low beta* (12–20 Hz), *high beta* (20–30 Hz), and *gamma* (30–45 Hz). Within each frequency band, the corresponding power *P*_*i*_ associated with *f*_*i*_ is calculated as in (35). Local power maxima are then identified to represent the dominant oscillatory activity in the cluster for a given event.

To investigate the relationship between oscillatory activity and memory encoding, a logistic regression model was trained using the dominant power values as predictors and the type of event (encoding of an arbitrary letter vs. fixation events) as the dependent variable for each individual subject. We adopted a liberal screening threshold of *p <* 0.25, following the recommendation of Hosmer & Lemeshow (Hosmer et al., 2013),

who argue that a conventional *p <* 0.05 threshold at this stage “often fails to identify variables known to be important,” particularly in small or noisy datasets. Given the inherent constraints of EEG data acquisition and the sample size, this choice was made to avoid prematurely discarding potentially relevant predictors. We emphasize that the present study is exploratory and hypothesis-generating in nature; our goal is not to make definitive inferential claims, but to identify candidate frequency-band correlates of memory encoding that merit investigation in larger, confirmatory studies. Under this screening criterion, the analysis revealed target p-values for high-beta and theta components in the frontal midline cluster for several individuals, indicating a plausible correlation between these frequency bands and memory encoding. The results are summarized in Fig. 7, where markers of the same color belong to a particular subject and p-values below 0.25 indicate candidate associations.

**Fig. 7.**
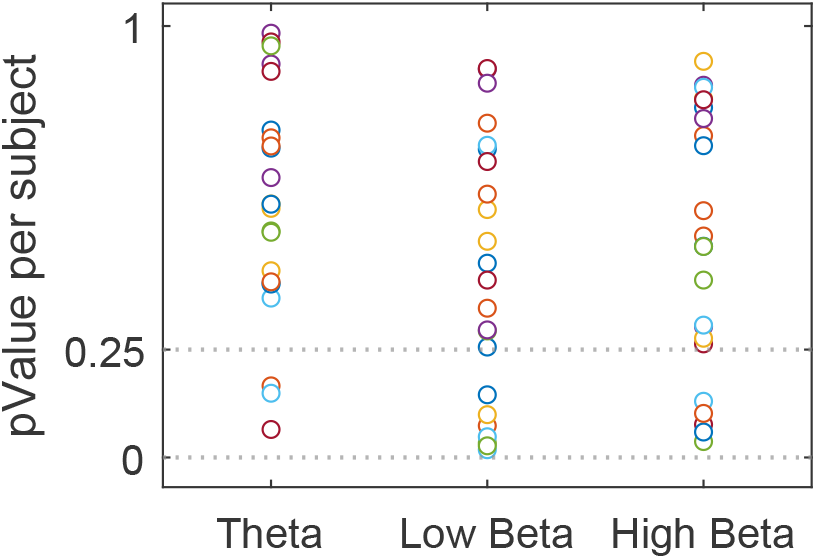
Frequency band against pValues from the logistic regression analysis on 23 subjects. Markers of the same color belong to a particular subject. The dashed lines indicate pValues below 0.25. 3 subjects are below this threshold in the Theta band, 6 Subjects in the Low-Beta, and 5 for the High Beta band.

We are interested in a possible correlation between theta power and memory load in the frontal midline cluster. For runs with a minimum of 5 encoded letters, total theta power per subject is calculated. Subjects are grouped in theta power percentiles (90th, 70th, 50th, and 30th), and we calculate Pearson correlation coefficients for theta power with respect to the number of encoded items. This analysis is included to replicate the distributional approach used in Onton et al. (2005), were it is found that memory load increases in theta power were driven primarily by a relatively small proportion of trials with particularly strong theta activity. The percentile analysis is included to investigate this observation with the RO-eDMD framework and is reported in Fig. 8. Despite the modest value of most coefficients, the analysis indicates that the more theta power, the more correlation: note how for almost every subject the 90th percentile is clearly the most correlated. Overall correlation coefficients and associated p-values are shown in Fig. 9, suggesting the effect, where present, may be driven by a subset of subjects rather than being consistent across the sample.

**Fig. 8.**
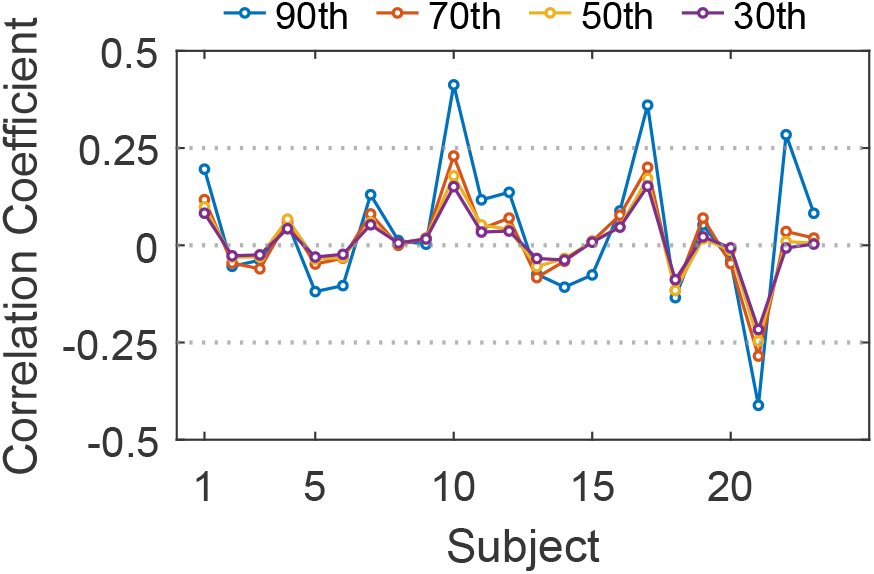
Subject versus Pearson correlation coefficient at 90th, 70th, 50th, and 30th of the theta power distribution. For most individuals, the runs with the most theta power present the most correlation

**Fig. 9.**
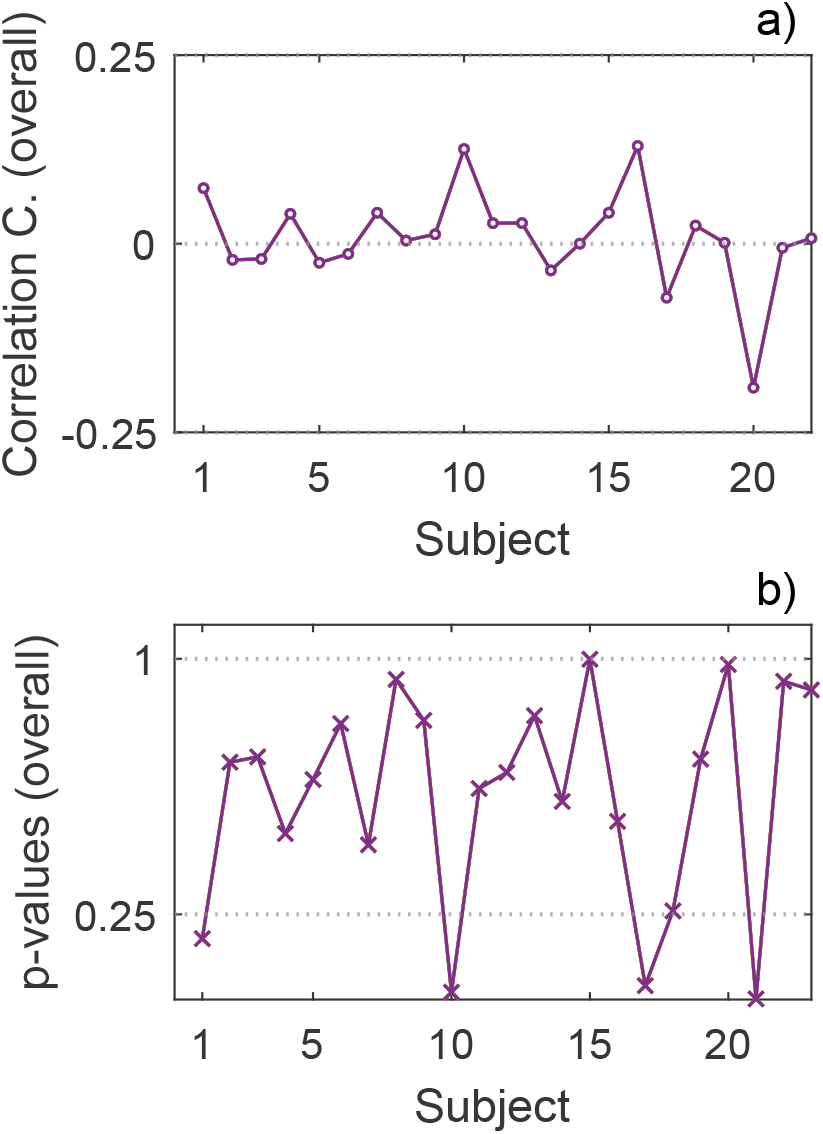
a) subject versus Pearson correlation coefficient. b) associated p-values

Our analysis replicates several key findings from (Onton et al., 2005) while also revealing important differences: Consistent with the original study, we observed a correlation between theta-band oscillatory power and memory encoding events in the frontal midline cluster. Our logistic regression analysis identified theta activity as a candidate predictor of encoding events for several subjects (Fig. 6). Our percentile-based correlation analysis (Fig. 7) confirmed the relationship between theta power and memory load stablished in Onton et al. (2005). Specifically, we found that the 90th percentile of theta power distribution showed the strongest correlation with memory load across most subjects, mirroring Onton et al.’s observation that memory-load-related increases in theta power were driven primarily by a relatively small proportion of high-theta trials (Onton et al., 2005).

A notable difference between our results and the original study is the frequency range of beta activity. Onton et al.’s study emphasized low-beta activity (12-15 Hz) as a consistent feature of frontal midline dynamics during memory encoding. By contrast, our logistic regression analysis identified correlations with activity in the high-beta range (20-30 Hz) rather than low-beta for several subjects (Fig. 6). We propose some possible explanations for this discrepancy:

- The methodological differences in frequency decomposition may account for this divergence. Onton et al.’s study employed Independent Component Analysis (ICA) to isolate fm*θ* components, followed by time-frequency analysis using event-related spectral perturbations (ERSPs) with a moving-window fast Fourier transform (FFT). Their low-beta band was defined as 12-16 Hz, and they noted that this activity sometimes appeared as harmonic energy from sharp-peaked theta wave trains. By contrast, the RO-eDMD approach we present extracts frequencies directly from the eigenvalues of the Koopman operator, which may provide a different spectral decomposition with different dominant frequencies, particularly for non-sinusoidal or transient waveforms.
- Differences in the selected electrodes may contribute to the disparity. Onton et al.’s study identified a frontal midline cluster (fm*θ*) consisting of ICA-based components. Our analysis, by contrast, used a subset of channels (Fz, FCz, Cz, CPz, AFz, F1, F2) based on the clustering reported in their study and anatomical correlations from (Koessler et al., 2009). While we aimed to capture the same frontal midline activity, our channel selection may have isolated different oscillatory dynamics.

## 6 Discussion

Our conclusions address both the methodological contribution of Reduced-Observable extended Dynamic Mode Decomposition (RO-eDMD) as a viable tool for neural data analysis and the neuroscientific insights it offers into a specific working memory paradigm in the frontal midline.

### Methodological Contribution and Empirical Validation

The primary claim of this work is that RO-eDMD over a dictionaryof RFFs provides a computationally efficient, interpretable framework for analyzing high-dimensional neural time series. This claim is directly supported by the empirical evidence presented in Section 5.1. Our comparison with standard DMD and a polynomial eDMD algorithm shows that RO-eDMD offers a significant advantage. The NRMSE of the RO-eDMD reconstruction is smaller than that of the polynomial eDMD at the tested SNRs, with a considerable reduction in computation time and a superior ability to recover the spectral content (see Fig. 5:d-f and Table 1) as well as the transient character of the tested signals. This establishes the RO-eDMD key strength: it offers an efficient compromise between the overly simplistic linear model of DMD and the computationally expensive, dictionary-dependent standard eDMD.

We find RO-eDMD over a random Fourier features dictionary particularly suitable for investigating transient, oscillatory patterns of neural activity associated with working memory, but the method is applicable to diverse, multidimensional neural recording modalities. Even with a modest nonlinear dictionary, the approach yields a useful decomposition and parallels findings previously obtained with PCA-based analysis (Onton et al., 2005). However, we acknowledge that the algorithm’s performance is sensitive to several parameter choices, including the dictionary size *D*, kernel bandwidth *γ*, rank truncation *r*, and the Hankel stacking parameter *h*. The sensitivity to noise, as shown by the increasing NRMSE at low SNRs, is a limitation that must be considered in future applications. A more systematic and data-driven parameter selection strategy is a crucial direction for future research.

### Neuroscientific Insights and Critical Interpretation

Our neuroscientific claim is that the method identified candidate frequency-band correlates of memory encoding in the frontal midline cluster in the data of several subjects. This claim is supported by the results of the logistic regression analysis (Fig. 7/5), which, under a liberal screening threshold of *p <* 0.25, revealed associations with high-beta and theta activity for several subjects. We must emphasize the exploratory nature of this analysis. The correlation threshold was chosen to avoid prematurely discarding potentially important variables (Hosmer et al., 2013) and is intended for generating hypotheses rather than making definitive inferential claims. Further confirmatory studies with larger sample sizes and more stringent statistical controls are necessary to validate these findings.

Our findings converge with those of (Onton et al., 2005) on the same dataset, particularly in replicating the relationship between theta power and memory load. The percentile-based correlation analysis (Fig. 8) mirrors Onton’s observation: the association is driven by a small proportion of trials with high theta power. This replication is significant because it demonstrates that the RO-eDMD algorithm, with a fundamentally different preprocessing and decomposition approach, can recover the same distributional phenomena, thereby lending empirical support to both the original finding and the viability of our method.

The discrepancy between our observed high-beta activity and the original study’s emphasis on low-beta (12-15 Hz) warrants a critical interpretation. While methodological differences in frequency decomposition and channel selection are plausible explanations, they also suggest that different analytical methods can highlight different aspects of the same neural signal. The Koopman operator’s eigenvalue decomposition is fundamentally different from the FFT-based power analysis used in (Onton et al., 2005).

### Overall Assessment

We consider that algorithmic-level descriptions (in the Marr levels sense (Marr, 2010)) of working memory can be enriched with nonlinear, dictionary-based approaches such as RO-eDMD. Our replication of a key effect in the EEG data provide an argument for the efficacy of RO-eDMD. We are also aware that the neuroscientific conclusions drawn from this initial application are tentative and require further validation. As decoding and prediction results become increasingly established, nonlinear methods like RO-eDMD provide an analysis avenue.

### Future directions

This study employed a relatively simple non-linear dictionary as a proof of concept. More sophisticated dictionaries may capture richer non-linear functional relations in the multidimensional data generated by neural recordings. As noted in (Logie et al., 2021), the advancement of working memory research requires consensus on ‘benchmark’ working memory phenomena that must be explainable by any valid model. Dictionary-based methods such as RO-eDMD provide important opportunities to achieve such benchmarking.

As our understanding of these methods matures, they may offer a deeper understanding of the computational mechanisms of cognition. Moreover, dictionary-based approaches (Logie et al., 2021) provide an important framework for evaluating competing models of memory by testing them simultaneously against empirical data. Dictionaries incorporating time-delay (Kamb et al., 2020) embeddings can prove valuable for investigating synchronization processes, phase dynamics, and causality relations (Brunton et al., 2021). Future work should also explore computational optimizations, including tensor network formulations (Lücke and Nüske, 2022), which could enable online analysis of neural datasets.

## Acknowledgements

The authors thank Prof. B. De Schutter for his invaluable feedback and assistance in revising the manuscript. We also acknowledge the support of the Delft Center for Systems and Control, Faculty of Mechanical Engineering, TU Delft.

## Declarations and Statements

All authors certify that they have no affiliations with or involvement in any organization or entity with any financial interest or non-financial interest in the subject matter or materials discussed in this manuscript.

All authors contributed to the manuscript design. Conception, material preparation, and analysis were performed by David Chavez-Huerta. Mohammad Khosravi edited and commented on current and previous versions of the manuscript. All authors read and approved the final manuscript.

## Appendix A

The evolving implementation of the analysis code is publicly available at: https://github.com/DchavezGH/ft_edmdanalysis.

The dataset used to generate the results supporting this manuscript is available at: https://github.com/DchavezGH/RO-eDMD.

## Footnotes

1 Note that rank reduction should allow for the preservation of the dynamics of interest in **Ψ**_*M*_ (**X**). Throughout this paper, we choose *r* to preserve up to 90% of the singular value energy, calculated as 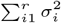, where *σ*_*i*_ denotes the singular values of **Ψ**_*M*_ (**X**), for *i* = 1, … , *m*, encoded in **Σ**_*f*_

2 In our algorithm, right eigenvectors *ξ*_*k*_ are projected on **V** to calculate **Φ**. Similarly, Koopman modes are calculated projecting left-eigenvectors on **V**^*\**^. This projections account for the rank reduction in 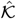.

## References

Brunton, S.L., Budišić, M., Kaiser, E., Kutz, J.N.: Modern Koopman theory for dynamical systems. arXiv preprint arXiv:2102.12086 (2021)

Brunton, B.W., Johnson, L.A., Ojemann, J.G., Kutz, J.N.: Extracting spatial–temporal coherent patterns in large-scale neural recordings using dynamic mode decomposition. Journal of Neuroscience Methods 258, 1–15 (2014)

Chavez-Huerta, D.: ft edmdanalysis. https://github.com/DchavezGH/ft_edmdanalysis (2026)

Colbrook, M.J.: The multiverse of dynamic mode decomposition algorithms. ArXiv abs/2312.00137 (2023)

Delorme, A.: Eeg is better left alone. Scientific Reports 13, 2372 (2023) 10.1038/s41598-023-27528-0

Hosmer, D.W., Lemeshow, S., Sturdivant, R.X.: Applied Logistic Regression, 3rd edn. John Wiley & Sons, New York (2013)

Kutz, J.N., Fu, X., Brunton, S.L.: Multiresolution dynamic mode decomposition. SIAM J. Appl. Dyn. Syst. 15, 713–735 (2015)

Kamb, M., Kaiser, E., Brunton, S.L., Kutz, J.N.: Time-delay observables for koopman: Theory and applications. SIAM Journal on Applied Dynamical Systems 19(2), 886–917 (2020)

Koessler, L., Maillard, L., Benhadid, A., Vignal, J.P., Felblinger, J., Vespignani, H., Braun, M.: Automated cortical projection of eeg sensors: Anatomical correlation via the international 10–10 system. NeuroImage 46, 64–72 (2009)

Logie, R.H., Camos, V., Cowan, N.: The state of the science of working memory. Working memory: State of the science, 1–9 (2021)

Lücke, M., Nüske, F.: tgEDMD: Approximation of the Kolmogorov operator in tensor train format. Journal of Nonlinear Science 32(4), 44 (2022)

Marr, D.: Vision: A Computational Investigation Into the Human Representation and Processing of Visual Information. MIT Press, Cambridge, MA (2010)

Onton, J., Delorme, A., Makeig, S.: Frontal midline eeg dynamics during working memory. NeuroImage 27, 341–356 (2005)

Onton, J., Makeig, S., Delorme, A., Truong, D., Robbins, K.: Sternberg Working Memory. OpenNeuro (2022). 10.18112/openneuro.ds004117.v1.0.1

Rahimi, A., Recht, B.: Random features for large-scale kernel machines. Advances in Neural Information Processing Systems 20 (2007)

Rahimi, A., Recht, B.: Uniform approximation of functions with random bases. 2008 46th Annual Allerton Conference on Communication, Control, and Computing, 555–561 (2008)

Roux, F., Uhlhaas, P.J.: Working memory and neural oscillations: alpha–gamma versus theta–gamma codes for distinct wm information? Trends in cognitive sciences 18(1), 16–25 (2014)

Sutherland, D.J., Schneider, J.G.: On the error of random fourier features. ArXiv abs/1506.02785 (2015)

Tu, J.H., Rowley, C.W., Luchtenburg, D.M., Brunton, S.L., Kutz, J.N.: On dynamic mode decomposition: Theory and applications. ACM Journal of Computer Documentation 1, 391–421 (2013)

Williams, M.O., Kevrekidis, I.G., Rowley, C.W.: A data–driven approximation of the Koopman operator: Extending dynamic mode decomposition. Journal of Nonlinear Science 25, 1307–1346 (2014)

Williams, M.O., Rowley, C.W., Kevrekidis, I.G.: A kernel-based approach to data-driven koopman spectral analysis. arXiv: Dynamical Systems (2014)

